# A repressor co-regulatory network integrates iron acquisition to mycobacterial stress tolerance

**DOI:** 10.1101/2025.02.12.637824

**Authors:** Khushboo Mehta, Kajal, Harsh Goar, Bhanwar Bamniya, Dibyendu Sarkar

**Author notes:** Department of Medicine, Division of Hematology-Oncology UT Southwestern Medical Center, Dallas, TX 75235. Address correspondence to: Dibyendu Sarkar, CSIR-Institute of Microbial Technology.

## Abstract

Growing evidence suggests that controlled regulation of iron (Fe) uptake is essential for mycobacterial survival and proliferation in the host. In this study, we discovered the mechanism that links mycobacterial Fe acquisition to intra-bacterial redox environment via the *phoP* locus. Remarkably, regulated expression of the major Fe storage protein encoding gene *bfrB* determines bacterial tolerance to oxidative stress and intracellular survival of the pathogen. Transcriptomic analysis coupled with *in vivo* DNA binding studies uncover a distinct IdeR-independent regulation, which utilizes Lsr2 and virulence regulator PhoP to modulate transcriptional control of *bfrB* via recruitment of both the regulators. A striking inhibition of Lsr2 binding to the *bfrB* promoter in a *phoP*-KO mutant, attributable to Lsr2-PhoP protein-protein interaction, provides the most fundamental biological insight. Building on these results, we proposed a model suggesting how Lsr2-PhoP interaction (or lack thereof) contributes to repression and/or stress-specific activation of *bfrB* expression. Collectively, these results uncover a key mechanism linking Fe acquisition and oxidative stress response of mycobacteria, and have significant implications on the intracellular survival of the pathogen.

## Introduction

Like most living organisms, *M. tuberculosis* utilizes Fe as a cofactor of important enzymes that are part of redox reactions and other essential cellular functions (1-3). As Fe level associates closely with oxidative stress for aerobic organisms, its deficiency often leads to lowered activity of heme-containing enzymes that offer protection against ROS. On the other hand, Fe overload due to dysregulated metabolism causes oxidative stress and excess Fe catalyses production of toxic oxygen radicals (4). To protect from Fe toxicity, *M. tuberculosis* has evolved regulatory control steps to maintain free intracellular Fe, absence of which sensitises the bacilli to macrophage killing and renders it growth attenuated in animal models (5-7). These results underscore that precise regulation of Fe uptake and homeostatic control on maintenance of intra-bacterial Fe remain essential for mycobacterial pathogenicity. Because of lower solubility at neutral pH, free Fe is not readily available within the host as it remains bound to high-affinity iron binding proteins. Thus, free Fe is present at a much lower concentration compared to what is necessary to support growth of the pathogenic bacilli under normal conditions. To counteract Fe limitation, *M. tuberculosis* secretes siderophores (mycobactins) which sequester Fe (III) and make it available to the bacteria via specialized Fe-siderophore transporters (8,9). In keeping with these, mycobacteria which fail to produce mycobactin shows defective bacillary multiplication in macrophages (10).

Prokaryotes often control intracellular Fe level by regulating transcription of genes related to iron acquisition (9). In mycobacteria, cellular Fe level is controlled at the transcriptional level by the Fe-dependent regulator IdeR, which in complex with Fe (II) regulates expression of ≍ 30 genes involved in synthesis, export and import of siderophores. IdeR binds to promoter region overlapping transcriptional start site and/or the -10 sequence, thereby interfering with RNA polymerase access to the promoter (11). Consequently, *ideR*-depleted *M. tuberculosis* shows uncontrolled Fe uptake and deficient Fe-storage leading to virulence attenuation due to toxic Fe overload (6). However, we still do not understand how transcriptional control of Fe acquisition and/or Fe metabolism contributes to mycobacterial tolerance to oxidative stress.

*M. tuberculosis* synthesizes many redox-stress inducible proteins like multiple regulators, thioredoxins, and alkyl hydroxy-peroxidase reductases under acidic conditions of growth. It has been shown that WhiB3-dependent lipid biosynthesis acts as a reductive reservoir to control redox balance during hypoxia (12) and macrophage infection (13), arguing in favour of the fact that *M. tuberculosis* under low pH conditions encounter reductive stress. These events lead to accumulation of NADH, and NADPH (14), and mycobacteria show a reduced cytoplasmic potential under acidic pH (15). However, mycobacterial cells encountering reductive stress in a paradoxical situation undergo adaptive metabolic changes to oxidize NADH/NADPH, and often generates ROS. These events account for elevated ROS level in mycobacteria with variable tolerance to oxidative stress under acidic pH compared to neutral conditions (16).

Given the fact that Fe acquisition is linked to oxidative stress response (5,17), and mycobacterial tolerance to oxidative stress is impacted by the *phoP* locus (18,19), we wished to explore role of PhoP in Fe acquisition and/or maintenance of Fe homeostasis and its effect on bacterial oxidative stress response. With the results uncovering a so-far-unknown role of PhoP in mycobacterial Fe uptake (Fig. 1), we next probed functioning of the regulator on transcriptional control of Fe metabolism genes. Our results demonstrate that PhoP in a context-dependent manner is recruited within promoters of Fe metabolism genes and regulates their expression including expression of major Fe storage protein BfrB, ectopic expression of which in WT-H37Rv rescues intra-mycobacterial Fe level under oxidative stress and promotes intracellular survival in macrophages. These results account for elevated sensitivity of *phoP*-KO mutant to oxidative stress (18,19). Further, we showed that PhoP synergizes with Lsr2 while functioning as a transcriptional regulator, directly represses *bfrB* expression, and Lsr2-PhoP protein-protein interactions function to control stress-specific bacterioferritin expression. Collectively, these results underscore the functional significance of stress-specific control of *bfrB* expression, and uncover an integrated regulatory mechanism of iron acquisition and its interdependence with mycobacterial oxidative stress response.

**Fig. 1:**
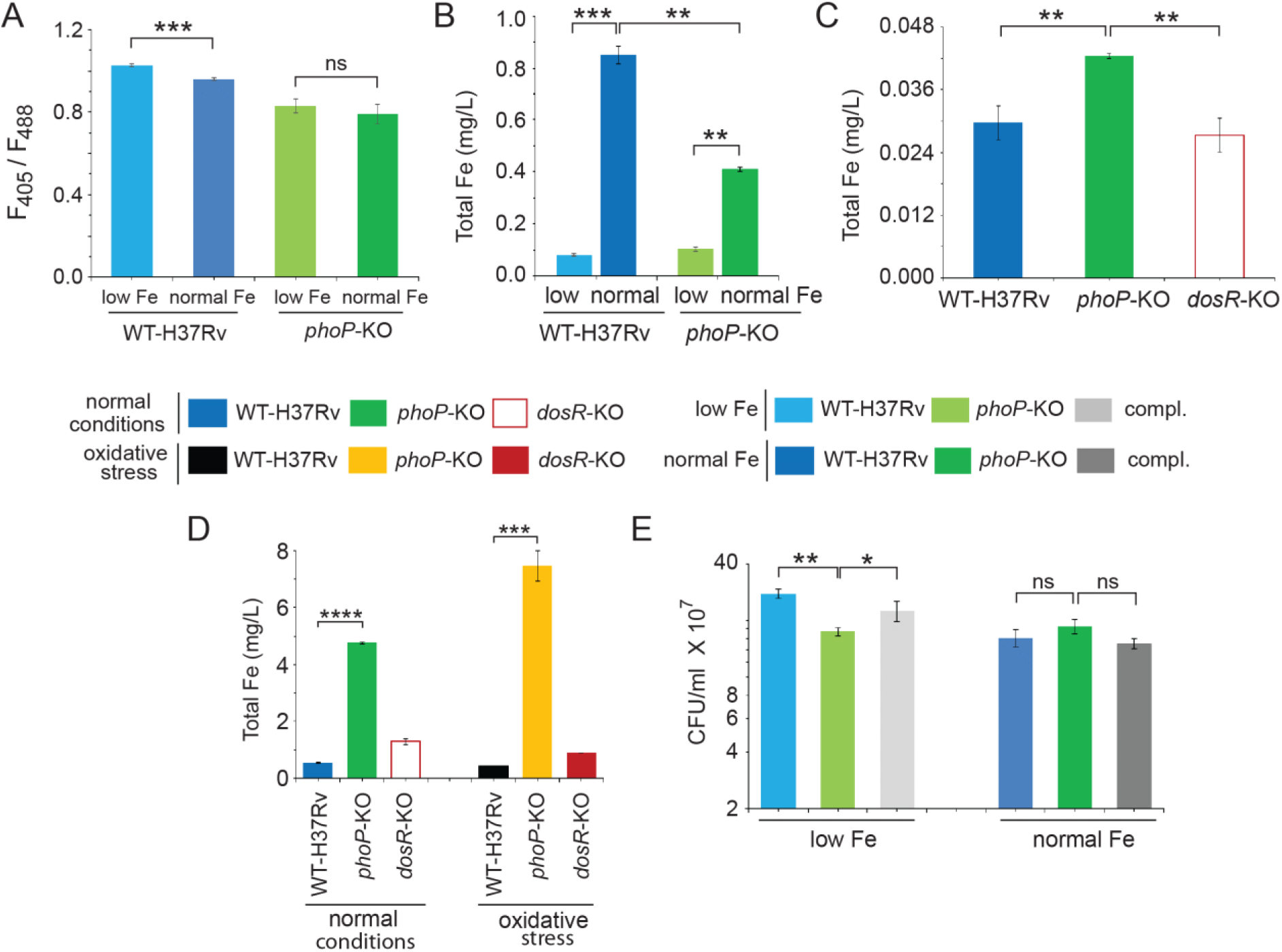
*phoP* links intra-bacteria Fe and mycobacterial redox poise. (A) Comparison of intra-mycobacterial mycothiol redox state of WT and *phoP*-KO grown either in low Fe or normal Fe conditions to examine whether *phoP* links intra-bacterial Fe to mycobacterial redox potential. This experiment utilized plasmid Mrx1-roGPF2 where mycoredoxin is fused to redox-sensitive GFP enabling real time measurement of mycothiol redox state by analysis of fluorescence emission at 510 nm after excitation of the samples at 405 and 488 nm, respectively (see methods). The results show average values from multiple replicates of experiments. (B) Intra-bacterial Fe levels were measured by IC-PMS for indicated strains grown under specific conditions as described in the Methods. Note that the difference in intra-bacterial Fe level in WT-H37Rv grown under normal conditions (*filled blue*) versus oxidative stress (*filled black*) is significant. (C) ICP-MS analysis of infected macrophage-derived *M. tuberculosis* harvested after 3-hour post infection showed a significantly elevated intra-bacterial Fe level of *phoP*-KO relative to WT-bacilli (see methods for details). As a control, *dosR*-KO strain under identical conditions showed comparable intra-bacterial Fe level as that of the WT bacilli. (D) WT-H37Rv and *phoP*-KO mutant, grown under normal conditions and oxidative stress, displayed a strikingly different level of intra-bacterial Fe content. However, identical comparison of intra-bacterial Fe level (compare *open red* versus *filled red* columns) shows insignificant variation for the *dosR*-KO mutant (used as a control strain) under identical experimental set up. The values represent average from multiple replicates of experiments. (E) To assess impact of available Fe in the growth media, WT-H37Rv, *phoP*-KO and the complemented mutant were grown in the presence of low or normal Fe concentration, and CFU values were enumerated from biological duplicates, each with multiple technical repeats.

## Results

### Mycobacterial Fe acquisition links intra-bacterial redox environment via the *phoP* locus

Previous studies suggest that iron (Fe) acquisition is linked to oxidative stress response of mycobacteria (5,17). To explore whether varying Fe concentration in the growth media influences intra-mycobacterial redox poise, we first preadapted, and then grew WT-H37Rv under low (0-2 µM) and normal (∼50 µM) Fe concentrations. As non-protein thiol MSH (mycothiol) is considered a substitute of glutathione in mycobacteria (20-22), we probed redox-dependent change in mycothiol redox state (Fig. 1A). Our assays used a redox-sensitive GFP (green-fluorescence protein)-based sensor fused to mycoredoxin (23), which enables measurement of mycothiol redox state in a real time scale. This probe was utilized to show that acidic pH inside the phagosome induces reductive stress for intra-phagosomal *M. tuberculosis* (24). Here, we found that WT-H37Rv grown under normal Fe conditions displays a more reduced environment relative to the bacilli grown under limiting Fe concentration (Fig. 1A). The difference is considered significant because even a minor alteration in fluorescence intensity ratios represent a significantly detectable change in the intracellular redox poise of mycobacteria (13,24-26). In fact, using this assay we recently showed that under oxidative stress WT-H37Rv displayed a significant increase in mycothiol redox state compared to unstressed mycobacteria (19).

As *M. tuberculosis* lacking a copy of *phoP* remains considerably more sensitive to oxidative stress relative to WT-H37Rv (18,19), we also grew *phoP*-KO strain and compared intra-mycobacterial redox-poise of the mutant grown in the media supplemented with low versus normal Fe concentrations (Fig. 1A). The objective was to examine whether Fe availability in the growth media of mycobacteria requires the *phoP* locus to determine mycothiol redox state. Note that the *phoP*-KO mutant displays a more reduced environment relative to WT-H37Rv both under limiting as well as normal Fe availability. More importantly, the mutant unlike WT-bacilli failed to show a differential redox environment when grown in the presence of low versus normal Fe concentration, indicating that mycobacterial Fe acquisition is linked to the *phoP* locus.

We next utilized Inductively Coupled Plasma Mass Spectrometry (ICP-MS) to determine intra-bacterial Fe concentration of indicated mycobacterial strains, grown under limiting and normal Fe concentrations. Fig. 1B compares total intra-bacterial Fe content of WT-H37Rv and *phoP*-KO mutant. WT-H37Rv showed ∼ 8-fold higher intra-bacterial Fe level when grown under normal Fe-containing media relative to bacterial cells grown under limiting Fe concentration. However, *phoP*-KO mutant grown under normal Fe-containing media showed ∼ 3.8 -fold higher intra-bacterial Fe level relative to the mutant grown under limiting Fe concentration. These results further connect *phoP* locus to mycobacterial Fe acquisition. We conclude that PhoP regulates mycobacterial Fe acquisition and contributes to variation of redox poise depending on the Fe availability in the environment.

To investigate role of the *phoP* locus during *in vivo* Fe acquisition, THP-1 macrophages were infected with WT-H37Rv and *phoP*-KO mutant at MOI of 1:10. 3-hours post infection *M. tuberculosis* strains were extracted from infected macrophages, subjected to lysis using 0.1% SDS and 0.2% nitric acid solution, and lysates analysed by ICP-MS to determine intra-bacterial Fe level (see methods for details). Importantly, *phoP*-KO mutant showed a significantly higher intra-bacterial iron level compared to the WT-H37Rv (Fig. 1C). However, under identical conditions, *dosR*-KO mutant showed comparable intra-bacterial Fe level as that of the WT-H37Rv. These results allow us to conclude that *M. tuberculosis* PhoP plays a major role in Fe acquisition *in vivo*. Notably, these findings are consistent with our *in vitro* data showing importance of PhoP in Fe acquisition of *M. tuberculosis*.

To investigate the link between oxidative stress and intra-bacterial Fe level, we next grew WT and *phoP*-KO mutant both under normal conditions and oxidative stress, and compared the intra-bacterial Fe levels (Fig. 1D). Notably, under normal conditions, the *phoP*-KO mutant showed ∼4-fold higher intra-bacterial Fe level relative to the WT-bacilli. However, we measured ∼7-fold higher intra-bacterial Fe level in the *phoP*-KO mutant relative to WT bacilli under oxidative stress. As a control, *dosR*-KO under identical conditions displayed an insignificant variation in intra-bacterial Fe level. From these results, we conclude that (a) intra-bacterial Fe level is closely linked to mycobacterial oxidative stress response via the *phoP* locus, and (b) dysregulated Fe acquisition of *phoP*-KO mutant under oxidative stress most likely accounts for bacterial hypersensitivity to specific stress conditions.

As *phoP* locus is linked to Fe acquisition of mycobacteria, we next counted viable bacterial colonies of WT and *phoP*-KO mutant under limiting versus normal Fe concentration, and enumerated the CFU values (Fig. 1E). Notably, *phoP*-KO strain shows a significantly reduced growth under limiting Fe concentration compared to WT-H37Rv, and complementation by stably expressing a copy of the *phoP* gene in the mutant restored CFU counts to the WT level. However, both WT-H37Rv and *phoP*-KO mutant displayed a comparable CFU in media containing normal Fe level. These results, for the first time, demonstrate that virulence regulator PhoP controls mycobacterial Fe acquisition.

### Complex transcriptional control of Fe metabolism genes requires the *phoP* locus

With the results identifying a new role of *phoP* locus on mycobacterial Fe acquisition, we next compared transcriptomic profile of WT-H37Rv and *phoP*-KO strain, grown under limiting or normal Fe concentration (Fig. 2). Our RT-qPCR results highlight that under limiting Fe conditions, two major genes related to Fe metabolism (*bfrA* and *irtA*) showed an elevated expression in the *phoP*-KO mutant relative to WT-bacilli (Fig. 2A). Complementation of the mutant by expressing a stable copy of *phoP* restored expression of these genes to the WT-level. These results suggest that under limiting Fe conditions, PhoP functions as a repressor of *bfrA* and *irtA*. In contrast, under normal Fe conditions, we did not observe a PhoP-dependent repression of *bfrA* expression (Fig. 2B). However, PhoP was found to repress *irtA* and 2 genes (*irtB and mbtB*) related to Fe metabolism displayed significant down-regulation in the *phoP*-KO mutant relative to WT-bacilli, suggesting PhoP-dependent activation of these genes under normal Fe concentrations. Note that we have previously shown reproducibly higher expression of *phoP* in the complemented mutant (relative to the WT bacilli) both under normal and low pH conditions of growth (27,28). These results possibly explain elevated mRNA levels of a few representative genes in the complemented mutant relative to WT-H37Rv. The oligonucleotides used in RT-qPCR experiments reported in this study are listed in Table S1.

**Fig. 2:**
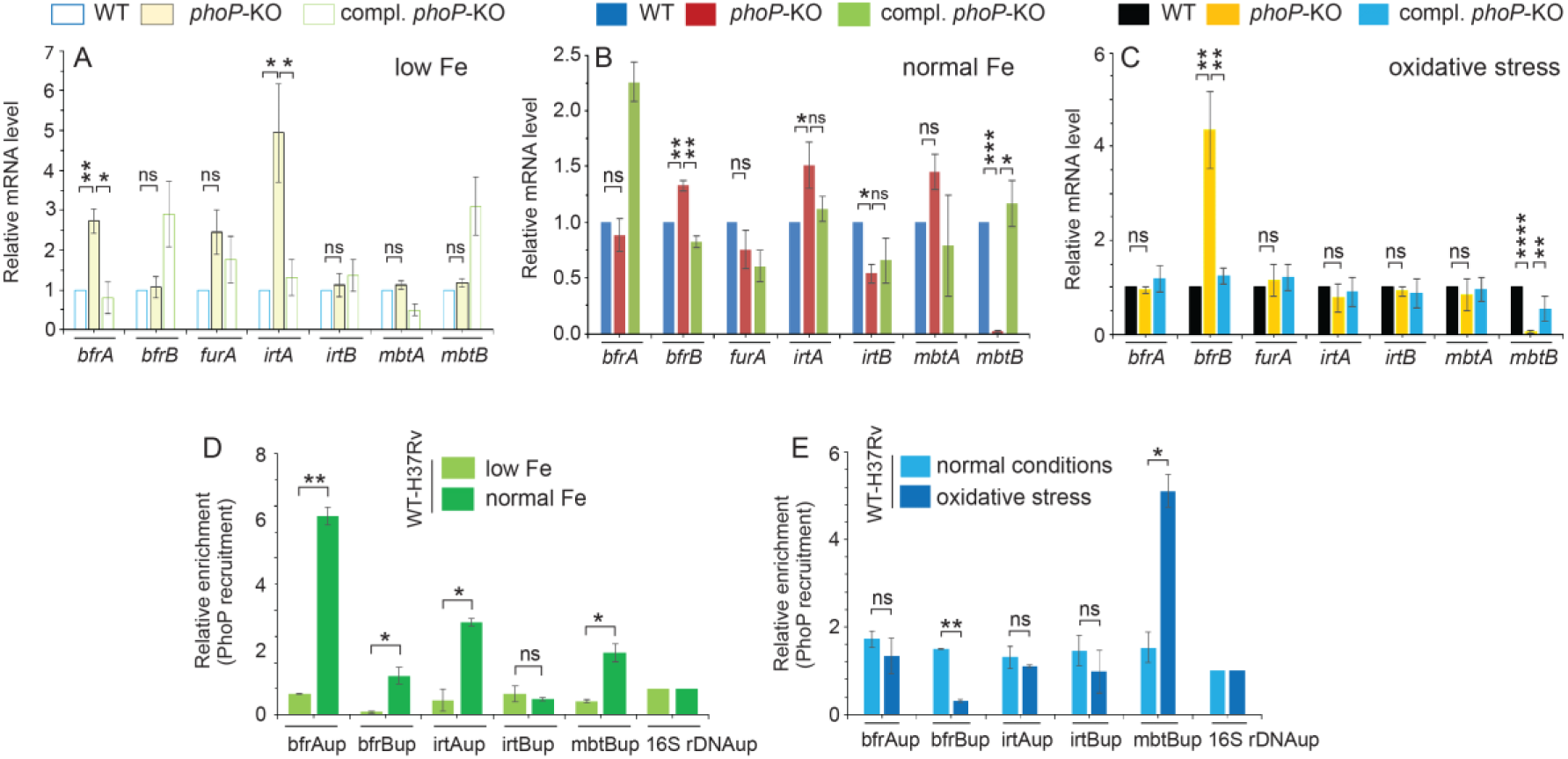
Expression of genes related to Fe metabolism require the *phoP* locus. Gene expression for representative mycobacterial genes was examined in WT-H37Rv, *phoP*-KO, and the complemented strain, grown under (A) limiting Fe, (B) normal Fe and (C) oxidative stress conditions using RT-qPCR as described in the methods. The data show plots of average values from multiple independent data sets. (D-E) To assess and compare context-dependent *in vivo* recruitment of PhoP within its target promoters, ChIP was carried out using anti-PhoP antibody, followed by qPCR using IP samples from WT-H37Rv grown under low Fe, normal Fe conditions and under oxidative stress (see methods for details). To determine fold enrichment, each data point was compared with the corresponding IP sample without adding antibody. The experiments were carried out from multiple biological replicates.

We next examined expression of these genes in mycobacterial strains grown under oxidative stress (Fig. 2C). The RT-qPCR results suggest PhoP-dependent and stress-specific repression of *bfrB* expression (compare Figs. 2B and 2C). In contrast, the two other PhoP activated genes (*irtA* and *irtB*) under normal conditions, remained unaffected by the regulator under oxidative stress. However, consistent with the mRNA levels under normal conditions, we noted a strong down regulation of *mbtB* in the *phoP*-KO mutant under oxidative stress. Collectively, PhoP-dependent contrasting regulation of *bfrB* expression under normal conditions versus oxidative stress (compare Figs. 2B and 2C) appears to function as a major determinant of mycobacterial oxidative stress response. Given the fact that *phoP*-KO, in one hand, shows an elevated level of *bfrB* under oxidative stress (relative to WT bacteria), and on the other hand, remains hyper-sensitive to oxidative stress, together these results suggest that precisely regulated *bfrB* expression accounts for mycobacterial tolerance to oxidative stress.

To investigate whether this regulatory effect is due to PhoP binding to it target sites, we assessed *in vivo* binding of the regulator by chromatin immunoprecipitation (ChIP) assays (Figs. 2D-E). In this experiment, cross-linked DNA-protein complexes of mycobacterial cells were grown under indicated conditions, fragmented to average size ≍500 bp by shearing, and real-time qPCR was performed using IP DNA and promoter-specific primer pairs (see supplemental Table S2) relative to sample without antibody as a control (mock). Our qPCR data demonstrate effective recruitment of PhoP within the target promoters. Notably, we observed Fe-concentration-dependent recruitment of PhoP within the upstream promoter regions of *bfrA*, *bfrB*, *irtA* and *mbtB* (Fig. 2D). These findings clearly align with PhoP-dependent regulation of these genes and functioning of the regulator in mycobacterial Fe acquisition. Furthermore, in agreement with the regulatory effect under oxidative stress, we observed stress-specific recruitment or lack thereof of PhoP within the upstream regulatory regions of *bfrB* and *mbtB*, respectively (Fig. 2E). From these results, we conclude that PhoP binds to the promoters of Fe metabolism genes in a context-sensitive manner.

### Ectopic expression of *bfrB* promotes bacterial tolerance to oxidative stress and improves intracellular survival

To explore the effect of PhoP-dependent over-expression of genes, we utilized an integrative expression plasmid pSTKi (29) and independently over-expressed *bfrA*, *bfrB* and *irtA* in WT-H37Rv. The oligonucleotides and plasmids used for amplification and cloning reported in this study are listed in Tables S3 and S4, respectively. Fig. S1A shows fold over-expression of genes in WT-H37Rv compared to the empty vector control. We grew these strains and compared intra-bacterial Fe concentration under normal conditions and oxidative stress (Fig. 3A). Although we observed a comparable level of intra-bacterial Fe under normal conditions of growth, a significantly higher Fe level under oxidative stress (relative to normal conditions) was apparent in these three strains relative to the empty vector control. We next measured tolerance of these strains to oxidative stress by determining the number of viable colonies (Fig. 3B). There was no significant change in CFU counts between WT-H37Rv strains overexpressing *bfrA* and *irtA* (WT::*bfrA* and WT::*irtA*) compared to WT-H37Rv containing an empty vector control. However, the *bfrB* over-expressing construct (WT::*bfrB*) displayed a significantly improved stress tolerance relative to WT-H37Rv. These results suggest that elevated intra-bacterial Fe level under oxidative stress is not the sole determinant of improved intracellular survival of mycobacteria. In fact, higher level of intra-bacterial Fe could be toxic and bactericidal excepting in case of *bfrB* over-expression, which effectively lowers mycobacterial sensitivity to oxidative stress. These results fit well with, and explain the previously puzzling observation that *M. tuberculosis* lacking a copy of the *phoP* gene is considerably more sensitive to oxidative stress relative to WT-H37Rv (18,19).

**Fig. 3:**
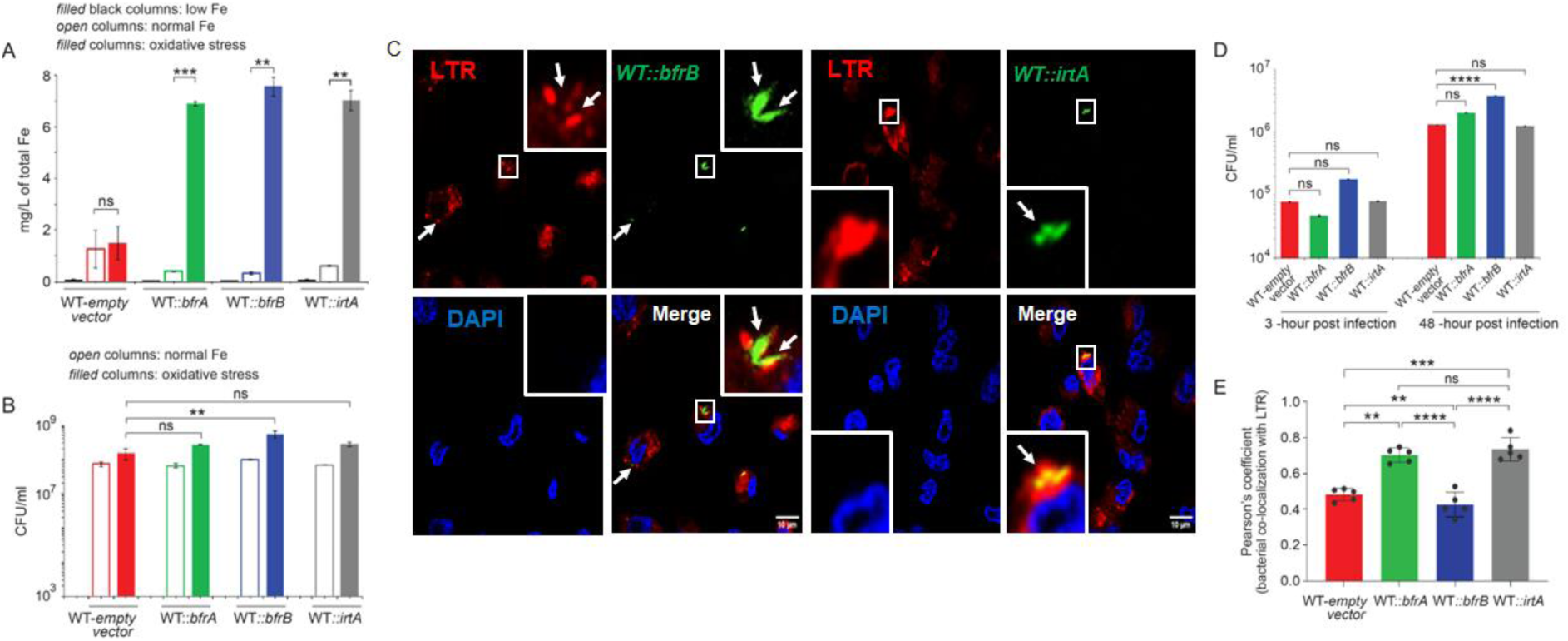
*bfrB* over-expression promotes intracellular survival of mycobacteria. (A) Intra-bacterial Fe levels were measured by IC-PMS for indicated over-expression strains, constructed by cloning the ORFs (see methods for further details), grown under limiting Fe, normal Fe and oxidative stress as described in the Materials and methods. The values represent average from multiple independent experiments. (B) To compare relative stress tolerance, the mycobacterial strains were next grown both under normal conditions and under oxidative stress. All four indicated strains displayed largely comparable growth under normal conditions without any significant difference. Further, similar to WT-H37Rv harbouring the empty vector, WT*::irtA* and WT*::bfrA* showed a comparable sensitivity to oxidative stress. However, WT*::bfrB* over-expressing bfrB displayed a significantly better growth as measured by the corresponding CFU values. The growth experiments were performed in multiple replicates. (C) To examine contribution of over-expression of specific mycobacterial genes on intracellular survival of the bacteria, THP-1 macrophages were infected with indicated bacterial strains. Mycobacteria and host cells were stained with phenolic auramine solution, and LysoTracker respectively. Host cell nuclei were made visible by Hoechst dye. Three fluorescence signals (Mycobacterial strains: green; lysosomes: red and host nuclei: blue) and their merge are displayed by confocal images (scale bar: 10 µm). (D) To examine mycobacterial survival in cellular models, THP-1 macrophages were infected with indicated mycobacterial strains, and 3-hour and 48-hour post infection intracellular bacterial CFU were enumerated. The results show average values from multiple biological replicates. (E) The data present Pearson’s correlation coefficient of images displaying internalized auramine-labelled mycobacteria and Lysotracker red marker in macrophages, and were evaluated using image-processing software NIS elements (Nikon). Five data points around the averages were shown, and average value with standard deviations were obtained from multiple independent experiments.

To investigate whether over-expression of these genes influence intracellular survival, we performed THP-1 macrophage infection experiments (Fig. 3C). Although WT-H37Rv carrying an empty vector could effectively inhibit phagosome-lysosome fusion, the two over-expressing H37Rv strains WT*::bfrA* (Fig. S1B) and WT*::irtA* (Fig. 3C) underwent phagosome-lysosome fusion, suggesting increased trafficking of the bacterial strains to lysosomes. In contrast, *bfrB* over-expressing H37Rv effectively inhibited phagosome -lysosome fusion (Fig. 3C). Notably, a comparable CFU values were determined for all of these strains 3-hr post infection (Fig. 3D). However, 48-hr post infection only *bfrB* over-expressing construct showed a significantly improved growth (∼3.7-fold) relative to WT-H37Rv. In contrast, under identical conditions both WT*::bfrA* and WT*::irtA* displayed an insignificant difference in growth within macrophages compared to WT-H37Rv. Consistent with these results, Pearson’s plot suggests that *bfrB* over-expression, but not *bfrA* or *irtA* over-expression, plays a major role in inhibiting phagosome maturation (Fig. 3E). Together, these results suggest that *bfrB*, encoding the major Fe-storage protein of *M. tuberculosis*, is critically required for bacterial growth within macrophages.

Since *mbtB* expression was significantly down-regulated in a *phoP*-KO mutant both under normal conditions and oxidative stress (Figs. 2B-2C), we explored impact of *mbtB* on intracellular survival of mycobacteria. We constructed a *mbtB*-KD mutant using a CRISPRi-based approach (30) and relative gene expression was compared with WT-H37Rv to confirm the knock-down mutant (Fig. S2A). In THP-1 macrophage infection experiments, *mbtB*-KD mutant, similar to WT-H37Rv (see Fig. S1B) could effectively inhibit phagosome -lysosome fusion (Fig. 4A). As a control, a *bfrB*-KD mutant (see Fig. S2B), under identical conditions underwent phagolysosome fusion (Fig. 4B), suggesting increased trafficking of the bacterial strain to lysosomes. In keeping with these results, *mbtB*-KD mutant displayed a comparable intracellular growth relative to WT-bacteria, whereas under identical conditions *bfrB*-KD showed a growth defect within macrophages (Fig. 4C). These results are consistent with previously reported growth of *mbtB*-KO mutant and WT-H37Rv within THP-1 macrophages comparable up to 48-hour post infection (10). Further, determination of Pearson’s coefficient suggests that bacterial colocalization with LTR (Lysotracker Red) is significantly impacted for *bfrB*-KD, but not for *mbtB*-KD (Fig. 4D). From these results, we conclude that under the conditions examined, *bfrB* but not *mbtB*, is essential for intracellular survival of mycobacteria.

**Fig. 4:**
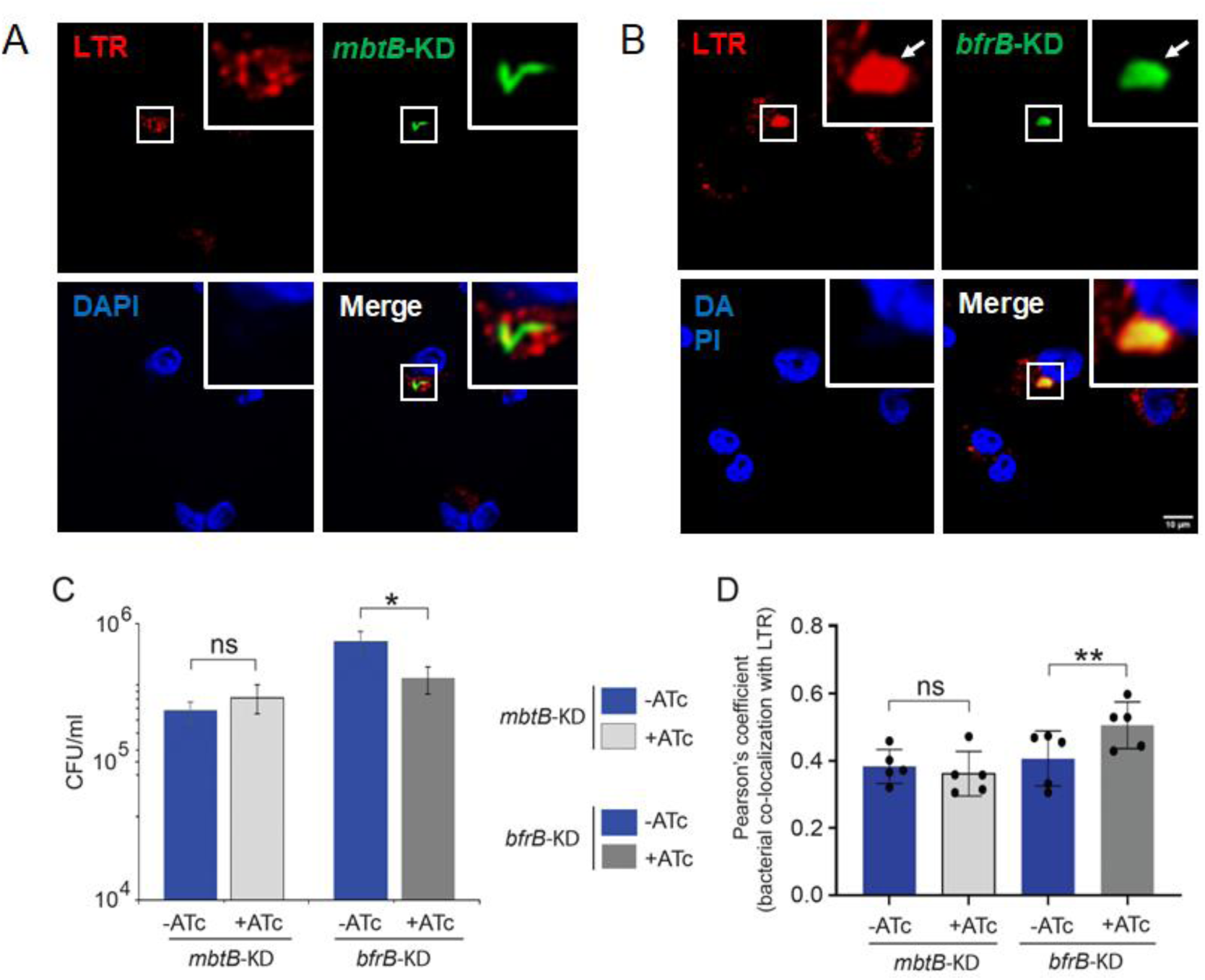
Knock-down mutant of *bfrB*, but not *mbtB*, displays intracellular growth defect of mycobacteria. **(A-B)** To investigate contribution of PhoP-controlled *mbtB* expression on intracellular survival of mycobacteria, THP-1 macrophages were infected with *mbtB*-KD mutant (details of the knock-down construct is described in the materials and methods). As a control, *bfrB*-KD mutant was used under identical conditions. Mycobacteria and host cells were stained with phenolic auramine solution, and LysoTracker respectively. Host cell nuclei were made visible by Hoechst dye. Three fluorescence signals (Mycobacterial strains: green; lysosomes: red and host nuclei: blue) and their merge are displayed by confocal images (scale bar: 10 µm). (C) To determine mycobacterial survival in cellular models, THP-1 macrophages were infected with indicated strains, and 48-hour post infection intracellular bacterial CFU were enumerated. The results show average values from multiple biological replicates. (D) The data present Pearson’s correlation coefficient of images displaying internalized auramine-labelled mycobacteria and Lysotracker red marker in macrophages. The results were evaluated and presented as described in the legend to Fig. 3E.

### Context-sensitive recruitment of regulators integrate Fe uptake and bacterial sensitivity to oxidative stress

Previous studies have shown that deletion of *M. tuberculosis* nucleoid associated protein (NAP) Lsr2, which functions as a gene silencing protein (31), significantly elevated transcription of *bfrB* (32). In keeping with these results, Rodriguez and co-workers showed that Lsr2 functions as a direct repressor of *bfrB* (33). Having unveiled a major role of PhoP in mycobacterial Fe uptake and transcription regulation of genes involved in Fe metabolism including a major regulatory impact on *bfrB* expression, we sought to ascertain how Lsr2 contributes to regulation of Fe metabolism. We adopted a CRISPRi-based approach (30) to construct *lsr2* knock-down mutant (see Methods for details). Determination of mRNA levels assessed expression of relevant genes in the knock-down constructs with respect to WT-H37Rv (Fig. S3A). *lsr2*-KD mutant displayed a significantly lowered expression of *lsr2*, and consistent with Lsr2-dependent repression of *fadD23* (31), *fadD23* level was significantly higher in the mutant strain relative to WT-H37Rv, confirming the identity of the mutant. Importantly, under normal conditions, we observed a significantly elevated expression of *bfrB* and *irtB* in the *lsr2*-KD mutant relative to WT-H37Rv, although expression of *bfrA*, and *irtA* remained largely unaltered (Fig. 5A). In an identical set-up, we also measured expression of these genes in a *phoP*-KD mutant (Fig. 5B). While *bfrA* expression remains unchanged, we noted down-regulation of *irtA*, *irtB* and a significant upregulation of *bfrB* in the *phoP*-KD mutant relative to WT-H37Rv. However, we noted that while Lsr2 represses *mbtB* expression, PhoP functions as an activator of *mbtB* (Fig. 5C). Fig. S3B confirms a significantly reduced *phoP* expression along with lowered expression of PhoP-activated genes in the *phoP*-KD mutant. Collectively, these data suggest that both Lsr2 and PhoP (a) participate as transcription regulators of Fe metabolism in mycobacteria and (b) appear to function as co-repressors of *bfrB* expression under normal conditions.

**Fig. 5:**
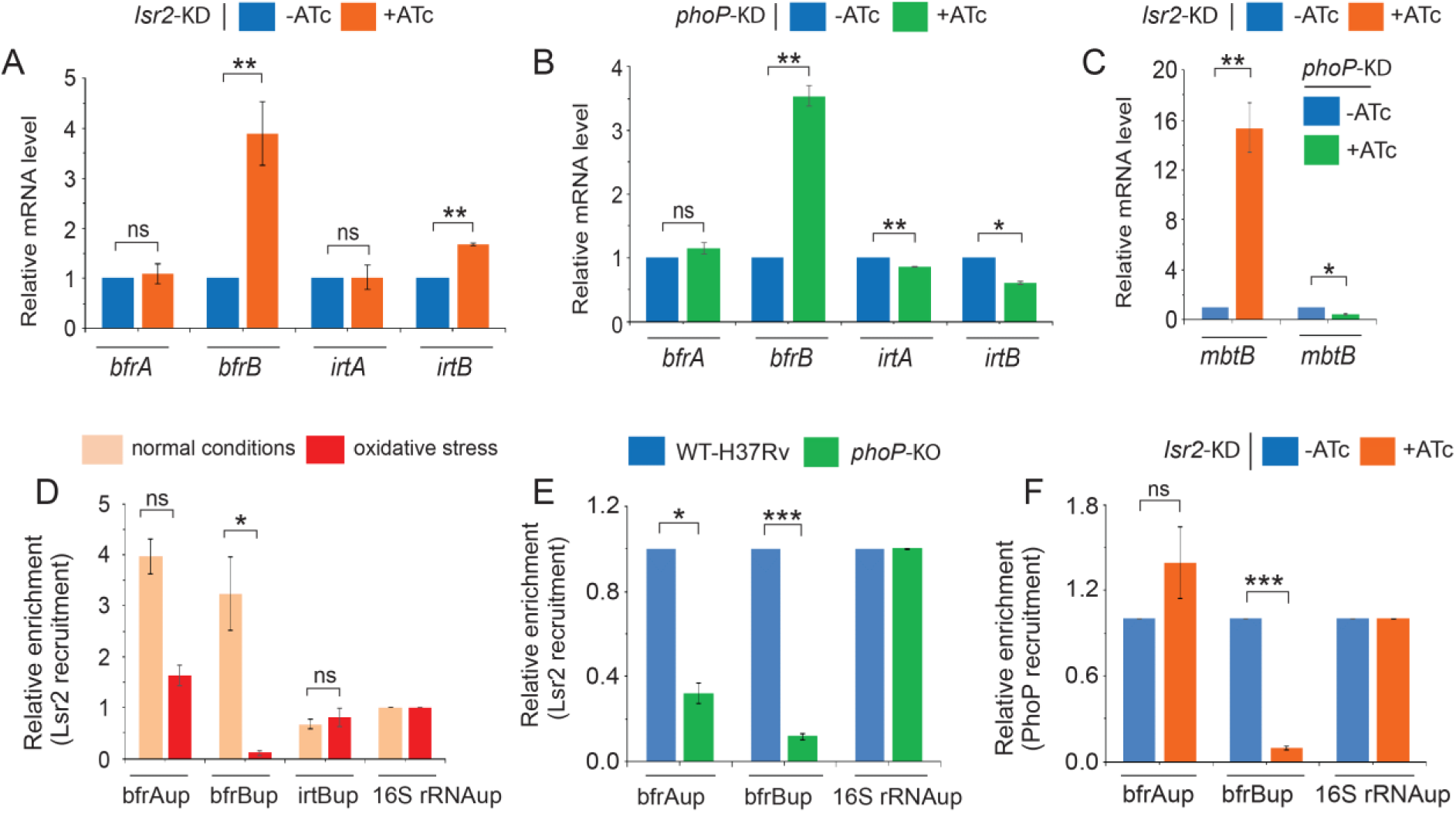
Lsr2 and PhoP function as co-repressors of *bfrB* expression. (A-B) mRNA levels of a few representative genes related to Fe metabolism were determined in (A) *lsr2*-KD and (B) *phoP*-KD mutants, respectively, grown under normal conditions of growth. Panel C compares *mbtB* expression in *lsr2*-KD and *phoP*-KD, grown under normal conditions. RT-qPCR measurements were performed as described in the Methods, and each value is an average of duplicate measurements originating from multiple biological replicates. (D-F) To assess and compare *in vivo* recruitment of Lsr2 (D-E) and PhoP (F) within its target promoters, ChIP was carried out using either anti-FLAG or anti-PhoP antibody, followed by qPCR using IP samples from WT-H37Rv (D), WT-H37Rv and *phoP*-KO (E) or WT-H37Rv and *lsr2*-KD (F), respectively. 3X-FLAG-tagged Lsr2 was expressed in WT-H37Rv and *phoP*-KO mutant using pNiT-1 expression system (see methods for details). To determine fold enrichment, each data point was compared with the corresponding IP sample without adding antibody. The experiments were carried out as multiple biological replicates, each with technical repeats.

To determine *in vivo* recruitment of Lsr2 within its target promoters, we next expressed 3X FLAG-tagged Lsr2 in WT-H37Rv (see methods) and performed ChIP experiments using anti-FLAG antibody (Fig. 5D). Thus, cross-linked DNA-protein complexes of growing bacterial cells were fragmented to an average size ≍500 bp by shearing, and DNA binding was assessed by qPCR. Table S2 lists the oligonucleotides used in ChIP-qPCR studies reported here. Our data demonstrate specific recruitment of Lsr2 compared to mock sample within its target promoters in WT-H37Rv. Together, the ChIP-qPCR experiments strikingly unravel that both Lsr2 and PhoP are recruited within the *bfrB* promoter under normal conditions, whereas both regulators fail to show effective recruitment under oxidative stress (compare Fig. 2E and Fig. 5D). Consistent with ChIP results, a PhoP binding site was identified ∼94-bp downstream of Lsr2 binding site (31) within the *bfrB* promoter (Fig. S3C).

To examine how the upstream regulatory region of *bfrB* accommodates both the repressors, we next compared *in vivo* DNA binding of Lsr2 in WT-H37Rv and a *phoP*-KO mutant (Fig. 5E). In this assay, we expressed a 3X-FLAG tagged Lsr2 in WT and mutant strain (see methods), and ChIP experiments were performed using anti-FLAG antibody. Likewise, to investigate a possible interdependence of the two regulators in DNA binding, we studied *in vivo* binding of PhoP in WT-H37Rv and a *lsr2*-KD mutant by ChIP assays using anti-PhoP antibody (Fig. 5F). Consistent with the regulation data, under normal conditions, both Lsr2 and PhoP showed specific recruitment within the promoters in WT-H37Rv (Figs. 5E and 5F, respectively). However, we observed a substantial reduction of (a) Lsr2 binding in *phoP*-KO mutant (Fig. 5E), and (b) PhoP binding in *lsr2*-KD mutant (Fig. 5F) within bfrBup relative to corresponding IP samples from WT-H37Rv, despite availability of comparable amounts of Lsr2 in WT- and *phoP*-KO mutant or PhoP in WT- and *lsr2*-KD mutant (Fig. S3D). Together, these results suggest that under normal conditions both regulators are recruited concomitantly within the target promoters, and recruitment of each of these requires the presence of the other.

### Probing Lsr2-PhoP interactions

We showed above that under normal conditions both Lsr2 and PhoP are recruited at proximal sites within the *bfrB* promoter. With the regulators functioning as co-repressors and showing an inter-dependent DNA binding, Lsr2-PhoP interaction was considered a possible mechanism to regulate *bfrB* expression. Thus, we probed Lsr2-PhoP interaction by using a previously reported protein fragment complementation (M-PFC) assay (34) (Figs. 6A-C). This approach uses bait and prey (two likely interacting mycobacterial proteins) as C-terminal fusions of two murine dihydrofolate reductase (mDHFR) fragments, that remain complementary to each other. When co-expressed in *M. smegmatis*, an interacting protein pair effectively reconstitutes a functionally active mDHFR, allowing bacterial growth in the presence of trimethoprim (TRIM). The plasmid pairs co-expressing corresponding protein pairs Lsr2/PhoP, IdeR/PhoP, and PhoP/PhoR (as a positive control) are described in the methods. Table S5 lists the oligonucleotides and plasmids used in this study. Lsr2 and IdeR were expressed from the episomal plasmid pUAB300 (Kan^R^), whereas *phoP* encoding ORF was expressed from the integrative plasmid, pUAB400 (Hyg^R^). In a different set, we have also expressed *phoP* from pUAB300, *lsr2* and *phoR* encoding ORFs from pUAB400, respectively. *M. smegmatis* co-transformants were screened on 7H10/Kan/Hyg, and sub-cultured in media containing antibiotics and 10 µg/ml TRIM. Although *M. smegmatis* co-expressing Lsr2 and PhoP grew well (Fig. 6A), strains harbouring empty vectors failed to grow in the presence of TRIM. Note that M-PFC assay using *M. smegmatis* co-expressing PhoP from episomal pUAB300 and Lsr2 from integrative pUAB400, respectively, displayed distinct bacterial growth on 7H10/TRIM plates (Fig. 6B). However, *M. smegmatis* co-expressing IdeR and PhoP, under identical conditions, failed to detect any bacterial growth on 7H10/TRIM plates (Fig. 6C). As a control, all of these strains showed normal growth in the absence of TRIM. From these results, we conclude specific protein-protein association between Lsr2 and PhoP.

**Fig. 6:**
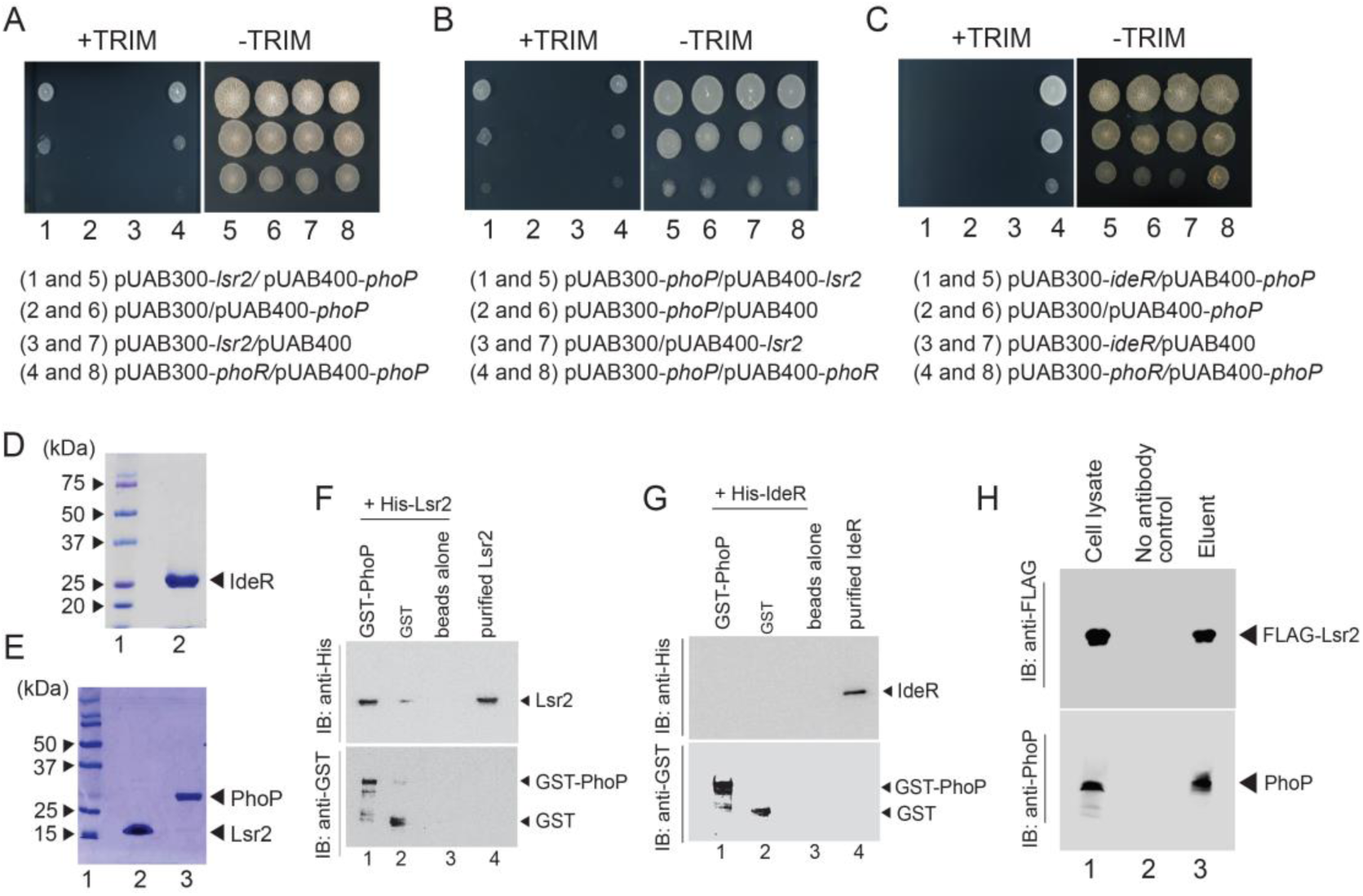
Lsr2 interacts with PhoP. (A-C) M-PFC experiments co-expressing *M. tuberculosis* Lsr2 and PhoP (A-B) or IdeR and PhoP (C) were used as screens using *M. smegmatis* as the surrogate host. Co-expression of plasmid pairs, as indicated under each panel, supporting *M. smegmatis* growth in the presence of TRIM is suggestive of specific interaction between Lsr2 and PhoP. Likewise, co-expression of a plasmid pair expressing IdeR and PhoP failed to support *M. smegmatis* growth in the presence of TRIM, suggesting lack of interaction between IdeR and PhoP. In each case, the three spots moving downwards in each panel represent spotting of cells at three different dilutions, namely undiluted, 10-fold and 100-fold dilutions, respectively. While co-expression of *phoP*/*phoR* in *M. smegmatis* displaying growth in presence of TRIM serves as a positive control, growth of all the strains in the absence of TRIM validates the assay. (D-E) For *in vitro* experiments, recombinant IdeR and Lsr2 carrying an N-terminal His_6_-tag (lane 2) were expressed and purified as described in the methods, analysed by SDS/polyacrylamide gel electrophoresis and visualized by Coomassie blue staining. As reference, molecular mass markers resolved alongside (lane 1) are indicated to the left. (F-G) To examine *in vitro* protein-protein interaction, crude extract expressing (F) His_6_-tagged Lsr2 or (G) His_6_-tagged IdeR was incubated with glutathione sepharose previously immobilized with GST-PhoP. Fractions of bound proteins (lane 1) were analyzed by Western blot using anti-His (upper panel) or anti-GST antibody (lower panel). Replicate experiments involved use of glutathione sepharose immobilized with GST alone (lane 2), or the resin alone (lane 3); lane 4 resolves purified Lsr2 or IdeR, respectively. (H) To probe interaction *in vivo*, 3X-FLAG-tagged Lsr2 was expressed in WT-H37Rv, and the cell lysate was immunoprecipitated with anti-PhoP antibody. Next, the IP material was probed by immunoblotting using anti-FLAG antibody (upper panel). Both Lsr2 and PhoP were detectable in the crude lysate (lane 1). Under identical conditions, samples from no antibody control (mock) (lane 2), remain undetectable by both anti-FLAG or anti-PhoP. However, eluent was detected for the presence of both proteins by anti-FLAG and anti-PhoP antibodies (lane 3), suggesting specific *in vivo* interaction between Lsr2 and PhoP.

We next examined Lsr2-PhoP interaction *in vitro*. To this end, recombinant IdeR and Lsr2 carrying an N-terminal His_6_-tag were cloned, expressed and purified as described in the methods (see Figs. 6D and 6E). Expression and purification of GST-PhoP has been described earlier (35). During *in vitro* pull-down assays GST-PhoP was immobilized to glutathione-Sepharose, followed by incubation with His_6_-tagged Lsr2. When proteins were eluted with 10 mM glutathione, both proteins could be detected in the same fraction (lane 1, Fig. 6F) in immuno-blot analysis. Under these conditions, GST-tag alone (lane 2) is at least 10 times less effective than the GST-PhoP (lane 1) in pulling down Lsr2 (based on the limits of detection in this assay). For only resin (lane 3) as a control, we could not detect both proteins in the same fraction. Together, these results suggest specific interaction between Lsr2 and PhoP. A similar pull-down experiment using His_6_-tagged IdeR and GST-tagged PhoP (Fig. 6G), could not detect both proteins in the same fraction (lane 1). These results are consistent with M-PFC data. From these results, we surmise that although IdeR does not interact with PhoP, Lsr2 interacts with PhoP.

To probe the interaction *in vivo*, we expressed 3X-FLAG-tagged Lsr2 in WT-H37Rv using pNiT-1 mycobacterial expression system (17,36) (see Methods for details). The whole cell lysate of WT-H37Rv was treated with anti-PhoP antibody, and the IP material was probed by Western blotting using anti-FLAG antibody. Expectedly, we could detect both Lsr2 and PhoP in the crude lysate (lane 1, Fig. 6H). However, samples from no antibody control (mock), under identical conditions, failed to detect FLAG-tagged Lsr2 (lane 2), suggesting specific interaction between Lsr2 and PhoP *in vivo*.

Because PhoP was previously shown to utilize its N-terminal domain to interact with other regulators (27,37,38), we next assessed role of PhoP domains as GST fusion constructs in Lsr2-PhoP interactions. Thus, truncated domains of Lsr2 and PhoP carrying an N-terminal GST-tag were cloned, expressed and purified as described in the methods (Figs. S4A and B). While previous studies have identified Lsr2 domain structure(39), we have shown that truncated domains of PhoP (PhoPN and PhoPC) remain functional for phosphorylation and DNA binding activity, respectively on their own (40). *In vitro* pull-down assays using N-domain of PhoP as a GST fusion (GST-PhoPN) with His-tagged Lsr2 displayed effective protein-protein interaction (lane 1. Fig. 7A) as that of full-length PhoP (see lane 1, Fig. 6F). However, under identical conditions, pull-down assays using His-tagged Lsr2 and GST-PhoPC suggest that C-domain of PhoP does not appear to contribute to Lsr2-PhoP interactions (lane 2). As controls, detectable signals were absent in case of GST alone (lane 3) or the resin alone (lane 4). To identify the corresponding interacting domains of Lsr2, we next probed Lsr2-PhoP interactions using His-tagged PhoP and GST-fusions of Lsr2 domains (Fig. 7B). Notably, GST-Lsr2N comprising Lsr2 residues 1-65 co-eluted with His-PhoP (lane 1). However, GST-Lsr2C comprising Lsr2 residues X-Y, under identical conditions, failed to co-elute with His-PhoP (lane 2). These results suggest specific interaction between Lsr2N and PhoP. Collectively, from these results we conclude that Lsr2-PhoP interaction is mediated by the N-domains of the corresponding regulators.

**Fig. 7:**
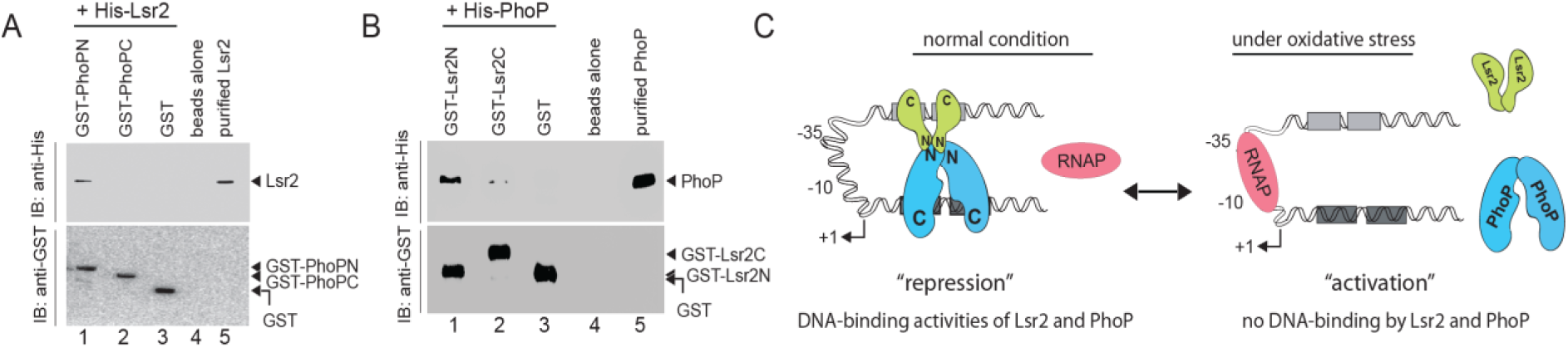
Lsr2-PhoP interaction is mediated by the corresponding N-terminal domains. (A) To examine *in vitro* interactions, crude extract expressing either His_6_-tagged Lsr2 was incubated with glutathione sepharose previously immobilized with GST-PhoPN (lane 1), GST-PhoPC (lane 2) and GST -tag alone (lane 3). Likewise, in panel (B), crude extract expressing either His_6_-tagged PhoP was incubated with glutathione sepharose previously immobilized with GST-Lsr2N (lane 1), GST-Lsr2C (lane 2) and GST -tag alone (lane 3). In both cases, fractions of bound proteins (lane 1) were analysed by Western blot using anti-His (upper panel) or anti-GST antibody (lower panel). Control experiments involved use of glutathione sepharose immobilized with the resin alone (lane 4); lane 5 resolved purified Lsr2 or PhoP, respectively. (C) Schematic summary of oxidative stress -dependent regulation of *bfrB* expression. In this model, we demonstrate that under normal conditions both Lsr2 and PhoP remain bound to their target sites preventing access for mycobacterial RNA polymerase (RNAP) to bind to the *bfrB* promoter and initiate transcription. Thus, gene expression is repressed under normal condition and additional stability of the higher order complex is attributable to Lsr2-PhoP protein-protein interaction. However, under oxidative stress both the regulators dissociate from their target sites allowing RNAP to bind to the *bfrB* promoter and initiate transcription.

## Discussion

The primary objective of this study was to understand how mycobacteria integrate Fe metabolism with oxidative stress response. First, we showed that WT-H37Rv shows a significantly different intra-bacteria redox poise when grown under limiting versus surplus Fe conditions (Fig. 1). The fact that *phoP*-KO mutant was considerably less tolerant to oxidative stress relative to WT-H37Rv led us to speculate that PhoP could be a major player connecting Fe metabolism and oxidative stress response. In line with the hypothesis, we were unable to detect a significant difference in mycothiol redox potential for a *phoP-KO* mutant of *M. tuberculosis*, grown under limiting versus surplus Fe conditions. Further, coupled with results showing significantly higher intra-bacterial Fe level for the macrophage-derived mutant bacteria (relative to WT-H37Rv), *in vitro* we noted considerably higher intra-bacterial Fe levels in the mutant relative to WT-H37Rv both under normal conditions and oxidative stress (Fig. 1). Together, these results unveil PhoP as a new player in the maintenance of mycobacterial Fe homeostasis.

Next, a transcriptomic approach to probe impact of PhoP on gene expression revealed PhoP-dependent striking repression of *bfrB* expression under oxidative stress (Fig. 2). Given the fact that *bfrB* encodes a major Fe storage protein(17,41,42) and *phoP*-KO mutant remains hypersensitive to oxidative stress (18,19), we speculated that regulation of *bfrB* expression accounts for mycobacterial oxidative stress response. This is consistent with ectopic expression of *bfrB*, but not *bfrA* or *irtA*, favouring bacterial growth under oxidative stress and in macrophages (Fig. 3). These results fit well with, and explain the previously reported observations by Pandey and Rodriguez that *M. tuberculosis* lacking a copy of the *bfrB* gene leads to Fe-mediated oxidative stress, and lowered bacterial pathogenicity relative to WT-H37Rv (5). It is noteworthy that *bfrB* over-expression either in *phoP*-KO under oxidative stress (Fig. 2) or ectopically in WT-H37Rv (Fig. 3) displays a contrasting growth phenotype both *in vitro* or in an intracellular milieu. These results suggest that in addition to regulation of *bfrB* expression, pleiotropic effects of PhoP affect global regulation of mycobacterial physiology.

*M. tuberculosis* lacking *bfrB* undergoes Fe-dependent oxidative stress, displaying elevated sensitivity to antibiotics and lowered pathogenicity (5). Consistent with these, a *bfrB*-KD mutant upon infection with macrophages readily undergoes phagolysosome fusion (Fig. 4). Paradoxically, ectopic expression of *bfrB* in an *ideR* mutant rescues Fe toxicity and complements bacterial survival in macrophages (6). In keeping with these results, we observed improved survival of *bfrB* over-expressing *M. tuberculosis in vitro* and in macrophages (Fig. 3). Collectively, these results highlight the importance of regulation of *bfrB* expression in mycobacteria. An attempt to identify mechanism of *bfrB* activation previously uncovered that IdeR binds to IdeR binding boxes within the promoter, and counters Lsr2, a histone-like protein that functions as a repressor of *bfrB (33),* suggesting a novel role of this regulator in Fe homeostasis. Although these results explain increased levels of *bfrB* in *lsr2*-deleted *M. tuberculosis* (32), the mechanism of regulation of *bfrB* under oxidative stress remained unknown. In this study, we show that *bfrB* is significantly repressed by PhoP under oxidative stress (Fig. 2). Consistent with this result, a PhoP binding site within the upstream regulatory region of *bfrB* was identified proximal to the Lsr2 binding site (Fig. S3C). Thus, we wished to explore how recruitment of two repressors within the same upstream regulatory region correlates with intricate control of *bfrB* expression.

A key question coming up in this study is whether Lsr2 communicates with PhoP *in vivo*. To investigate this, we compared *in vivo* Lsr2 recruitment at the target promoter(s) in WT-H37Rv and *phoP*-KO (Fig. 5E). Remarkably, Lsr2 recruitment within the *bfrB* promoter was significantly lower in a *phoP*-KO mutant compared to WT-H37Rv. Likewise, a significantly reduced PhoP recruitment at the *bfrB* promoter was observed in *lsr2*-KD mutant relative to WT-H37Rv (Fig. 5F). As simultaneous presence of Lsr2 and PhoP or lack thereof at the *bfrB* promoter, under normal conditions and oxidative stress, respectively was required for stress-specific *bfrB* expression (Figs. 2E and 5D), we explored Lsr2-PhoP protein-protein interaction. Our results suggest that Lsr2 interacts with PhoP (Fig. 6), and most likely Lsr2-PhoP interaction accounts for context-specific complex control of *bfrB* expression. Although, we cannot exclude the possibility that down-regulation of *bfrB* expression by Lsr2 and PhoP are unrelated to each other, we consider this an unnecessary complexity and therefore, we favour a more integrated view of the results considering that Lsr2-PhoP interaction controls stress-specific regulation of *bfrB* expression as depicted schematically in Fig. 7C. Thus, under normal conditions, Lsr2-PhoP interaction stabilizes higher-order DNA-protein structure, inhibits RNAP to bind to the *bfrB* promoter, leading to repression of *bfrB* expression. However, under oxidative stress, the regulators come off their respective cognate sites providing access to RNAP to initiate transcription, and thereby de-repress promoter activity. Our view on the working model retains the role of phosphorylation-coupled DNA binding of PhoP, as proposed in a model of low pH-inducible mycobacterial gene expression (28), and it focuses attention on the importance of N-terminal domain in context-sensitive regulation of gene expression. We also suggest that, based upon what is known about the response regulator family, it would not be surprising at all if this critical new role of the N-terminal domain of PhoP is a commonly shared basic feature of other members of the family.

Bacterioferritin expression in *M. tuberculosis* is most likely under precise regulation as survival of the pathogen under oxidative stress is essential for a successful infection. Given the fact that Lsr2 is capable of bridging distant DNA sequences (43), the finding that PhoP binding guides Lsr2 recruitment within *bfrB* upstream regulatory region, provides us with a fundamental biological insight into the context-dependent regulation of *bfrB* expression. In such a situation, Lsr2 might effectively function by binding to *bfrB* promoter already bound to PhoP. Thus, a complex interplay of Lsr2 and PhoP most likely fine tunes precise regulation of *bfrB* expression, and perhaps this is appropriate for the regulated expression of a gene which plays such a critical role in mycobacterial oxidative stress response and intracellular survival. It should be noted that the results reported here uncovering guided recruitment of a NAP by a response regulator as a functional partner, controlled by protein-protein interactions, represent an additional tier of regulation that have perhaps evolved to further regulate niche-dependent control of gene expression. Thus, a rather subtle intricate control of regulated *bfrB* expression during infection may be attributable to interplay of multiple regulators. In fact, earlier studies have shown a few pair of regulators like EspR/PhoP and CRP/PhoP interact functionally during transcriptional control of ESAT-6 secretion, and cAMP-inducible gene expression, respectively (27,44). It is conceivable that to fine tune niche-specific regulation of gene expression for critically important genes, this may be a preferred mechanism utilized by prokaryotes than has been previously acknowledged.

## Materials and methods

### Bacterial strains, media and growth conditions

*Escherichia coli* strains XL10 (Stratagene) and BL21(DE3) (Novagen) were used for cloning and protein over-expression, respectively. These were grown in Luria-Bertani (LB). *M. smegmatis* mc^2^155 was grown in Middlebrook 7H9 (m7H9) media or 7H10 agar containing with 0.2 % glycerol and 0.05% Tween-80. PhoPR-deleted *M. tuberculosis* (*phoP*-KO) and the complemented mutant have been described earlier (18). DosR-deleted *M. tuberculosis* (*dosR*-KO) was constructed by ‘mycobacterial recombineering’ as described (37). Wild-type *M. tuberculosis* H37Rv (WT-H37Rv), *phoP*-KO and *dosR*-KO or their derivatives were cultured at 37°C in m7H9 containing BSA, 0.2% dextrose and 0.85% NaCl (ADS, albumin-dextrose-sodium chloride). For bacterial growth under low Fe conditions, minimal medium containing 0.5% w/v asparagine, 0.8% w/v KH_2_PO_4_, 2% glycerol, 0.05% Tween-80 and 10% ADS, was used to culture mycobacterial strains. Chelex-100 (Bio-Rad) was used as per recommendations of the manufacturer to reduce trace metal concentrations within 0-2 µM as described previously (33). While this was considered as low Fe medium, normal Fe medium was generated by supplementing low Fe medium with 50 µM FeCl_3_. Transformation, selection of transformants, and subsequent growth studied using various mycobacterial strains were performed as described elsewhere (45). For growth under oxidative stress, mycobacteria were grown to OD_600_ ∼ 0.4-0.6, and further grown with 5 mM diamide for 2 hours. For CFU measurements, strains were inoculated into freshly prepared 7H9-ADS medium supplemented with 5 mM diamide (Sigma) at OD_600_ ∼ 0.05, and additional growth was allowed for 48 hrs at 37°C. To determine CFU values, cells were grown for 21 days on 7H11-OADS (oleic acid-ADS) agar plates. Where appropriate, kanamycin (kan), 20 µg/ml; hygromycin (hyg), 50 µg/ml; and ampicillin (amp), 50 µg/ml were used as antibiotics.

### Quantification of Fe content in bacterial cultures

Mycobacteria was grown to OD_600_ ∼ 0.8-1.0 and total intra-bacterial Fe concentrations were determined by ICP-MS. Cells were harvested, and washed with 1X PBS for three times. Finally, mycobacterial cells were lysed by boiling in 0.1% SDS and 0.2% HNO_3_ for 15 minutes in trace-element free 1.5 ml micro-centrifuge tubes. Following lysis, total volume of each sample was made up to 1 ml using MS grade water, samples were filtered through syringe filters, and analysed on ICP-MS (ThermoXcaliber II). To determine intra-bacterial Fe content in macrophage-derived mycobacteria, THP-1 cells were infected with WT-H37Rv and *phoP*-KO mutant of *M. tuberculosis* at an MOI of 1:10. 3 hours post infection, macrophages were lysed with 0.05% SDS (prepared in MS grade water) and centrifuged at 4000 rpm for 8 minutes. The bacterial pellet was subjected to lysis by boiling in 0.1% SDS and 0.2% nitric acid solution for 15 minutes in trace-element free 1.5 ml micro-centrifuge tubes. Volume was made up to 1 ml using MS grade water and samples filtered before analysis by ICP-MS.

### Cloning

To over-express mycobacterial genes *irtA* and *bfrA*, both ORFs were amplified using primer pairs FPirtA/RPirtA and FPbfrA/RPbfrA, respectively, cloned and expressed from the integrative vector pSTKi (29). Likewise, *bfrB* was amplified using FPbfrB/RPbfrB primer pair and cloned between BamHI and HindIII sites of pSTKi. Recombinant His_6_-tagged PhoP was cloned between NdeI and HindIII sites of pET-28b using the primer pair phoPstart/phoPstop (35) (see Table S3). Full-length IdeR, and Lsr2 were cloned between BamHI/HindIII sites of pET28a, and NdeI/HindIII sites of pET28b using primer pairs FPideRstart/RPideRstop and FPlsr2start/RPlsr2stop, respectively. To express *M. tuberculosis* PhoP in mycobacteria, the ORF with an N-terminal Flag-tag was cloned, and expressed from p19Kpro (46) as described previously (44). To express Flag-tagged IdeR and Lsr2, amplifications were carried out using primer pairs FPideRNiT/ RPideRNiT and FPm1lsr2/RPm1lsr2, and cloned between EcoRV/HindIII and HindIII/PacI sites of pNiT-1 (36), respectively. *dosR*-KO was complemented by stably expressing a copy of *dosR* using integrative vector pSTKi (29). *lsr2* carrying a carboxy-terminal FLAG tag was cloned in p19Kpro between HindIII and ClaI sites using primer pair FPm1Lsr2/ RPm2Lsr2 and expressed in *phoP*-KO mutant. To express truncated Lsr2 domains, plasmids pGEX-*lsr2N* (195 bp of the ORF comprising Lsr2 residues 1-65) and pGEX-*lsr2C* (180 bp of the ORF comprising residues 53-112) were constructed by amplifying the corresponding ORFs using PCR primer pairs FPlsr2N**/**RPlsr2N, and FPlsr2C**/** RPlsr2C, and cloning these amplicons between BamHI/XhoI and EcoRI/XhoI sites of pGEX 4T-1 (GE Healthcare), respectively. The resulting constructs GST-Lsr2N and GST-Lsr2C were expressed with an N-terminal GST-tag. Likewise, to express GST-tagged *phoPN* (encoding 423-bp of the N-terminal *phoP* ORF), and *phoPC* (encoding 321-bp of the C-terminal *phoP* ORF), PCR products were amplified and cloned in pGEX-4T-1 as described for pGEX-*phoP* (35). Each of the constructs was checked by DNA sequencing. Cloning and amplifications reported in this study required use of oligonucleotides and plasmids, which are listed in Tables S3 and S4, respectively.

### RNA isolation and Quantitative Real-time PCR

Mycobacterial cultures were grown to log phase under normal and indicated conditions and RNA samples were purified as described previously (27). Integrity of sample were assessed by gel electrophoresis. RNA samples were quantified by recording absorbance at 260 nm. Each RNA sample was treated with DNaseI for 30 minutes at room temperature to remove genomic DNA. Next, cDNA synthesis and RT-PCR conditions were followed as described previously (27) using Superscript III platinum-SYBR green kit (Invitrogen) and at least two independent RNA preparations. *M. tuberculosis rpoB* or 16S rDNA or *sigA* was used as endogenous controls. ΔΔC_T_ method was used to determine fold difference in gene expression as described elsewhere (47). Table S1 shows the oligonucleotide primers used in RT-qPCR studies reported here.

### Expression and purification of proteins

Recombinant PhoP, Lsr2 and IdeR proteins from *M. tuberculosis* were expressed as His_6_-tagged proteins and purified by Ni^+2^ -affinity chromatography as detailed elsewhere (35). Also, full-length PhoP, and truncated domains of Lsr2 and PhoP containing an N-terminal GST-tag were expressed and purified as described previously (35). The protein concentration was measured by Bradford reagent with BSA as the standard, and expressed in equivalent of protein monomers. The protein samples were stored at -80°C in 50 mM Tris-HCl, pH 7.5, 300 mM NaCl, and 10% glycerol.

### ChIP-qPCR

ChIP experiments were carried out as described (27). FLAG tagged IdeR, Lsr2 and PhoP were expressed in WT-H37Rv and immunoprecipitation (IP) was carried out using anti-PhoP or anti-FLAG antibody and protein A/G agarose beads (Pierce). To determine recruitment of the regulators, qPCR was performed in reaction buffers using appropriate dilutions of immunoprecipitated (IP) DNA, SYBR green mix (Invitrogen), specific primer pairs (0.2 µM; Table S2) that spanned promoter regions of interest and one unit of Taq DNA polymerase (Platinum Taq, Invitrogen). In all cases, results were compared with PCR signal from mock sample, which was obtained from an IP experiment without adding antibody. To examine non-specific recruitment, IP material from the samples were further analyzed using 16S rDNA or *rpoB*-specific oligonucleotide primers. In all cases analysis of melting curve was performed to verify amplification of a single product and qPCR reactions were performed in duplicate from two independent bacterial cultures.

### Macrophage Infections

In 6-well plates, THP-1 macrophages were seeded at a density of 1 x10^5^ cells/ml. Macrophages were infected with Titrated cultures of mycobacterial strains at a MOI of 10 bacteria per macrophage. *M. tuberculosis* H37Rv strains were stained with phenolic auramine, whereas macrophages were visualized by staining with Lyso-Tracker Red DND 99 (Invitrogen). Following fixation, cells were examined with a Nikon A1R confocal microscope. Macrophage infected with WT-H37Rv was used to optimize laser/detector parameters and IMARIS software (version 9.20) was used to process the digital images. Every image was subjected to a uniform intensity threshold, and percent bacterial co-localization was assessed by analysis of ∼ 50 infected cells of multiple fields from two independent biological repeats.

### Construction of knock-down mutants of *M. tuberculosis* H37Rv

Knock-down mutants of *bfrB* (*bfrB*-KD), *lsr2* (*lsr2*-KD), and *mbtB* (*mbtB*-KD) were created by utilizing a previously reported CRISPRi-based approach (28). This method uses target gene-specific guide RNAs (sgRNA) to inhibit gene expression by inducible expression of dcas9. To generate WT-H37Rv::dCas9, the integrative plasmid pRH2502 expressing *S. pyogenes* dCas9 was introduced in WT-H37Rv. Next, cloning of target gene-specific sgRNAs was carried out in pRH2521 using BbsI, and the oligonucleotide design ensured that the sgRNAs consisted of a 20-bp sequence complementary to the non-template strand. The clones were sequenced and introduced in *M. tuberculosis* harbouring pRH2502. In order to inhibit expression of specific genes by sgRNA and express dcas9, the bacterial cultures were grown in m7H9, supplemented with 0.2 % glycerol, 10 % OADC, 0.05 % Tween, 50 μg/ml hygromycin and 20 μg/ml kanamycin at 37°C. Anhydro-tetracycline (ATc) at a final concentration of 600 ng/ml was added at an interval of 48 hours, and cultures were grown for 4 days. RT-qPCR experiments using RNA samples of knock-down mutants examined expression of target genes. For the induced strains (in the presence of ATc) expressing sgRNAs targeting +234 to +253 (relative to *bfrB* translational start site), +56 to +75 sequences (relative to *lsr2* translational start site), and +56 to +75 sequences (relative to *mbtB* translational start site), we obtained approximately ∼98%, ∼75% and ∼70% reduction of *bfrB, lsr2,* and *mbtB* RNA levels, respectively, relative to the corresponding uninduced (in the absence of ATc) strains. The oligonucleotides and the plasmids utilized in knock-down experiments are shown in Table S3.

### Co-immunoprecipitation

To investigate protein-protein interactions *in vivo*, cell lysates of *M. tuberculosis* harbouring 3X-FLAG-tagged Lsr2 was immunoprecipitated with anti-PhoP antibody. To this end, ∼1.5 mg total protein and 50 µg of anti-PhoP antibody was incubated for overnight 4°C. An additional incubation of further 2 hours at 4°C was carried out in the presence of protein A/G agarose beads (∼20 µl) (Thermo Scientific). To confirm interactions, samples from no antibody control (mock) under identical conditions, and eluents from protein A agarose beads were analysed by immunoblotting.

### Mycobaterial protein fragment complementation (M-PFC) assays

*M. tuberculosis phoP* and *phoR* ORFs were amplified using primer pairs mphoPF/mphoPR and mphoRF/mphoRR, respectively. The amplicons were then cloned into the integration vector pUAB400(kanR) between the MfeI and HindIII sites and into the episomal plasmid pUAB300(hygR) between the PstI and HindII sites, respectively. Likewise, to express *M. tuberculosis phoP* from the episomal plasmid pUAB300, the ORF was amplified by the primer pair FPm2phoP/mphoPR, and cloned between BamHI and HindIII sites of pUAB300. Transformants (*M. smegmatis* mc^2^155 containing pUAB400-*phoP* or pUAB300-*phoP*) were further grown to prepare competent cells. Similarly, *M. tuberculosis phoR* was amplified by the primer pair FPm2phoR/mphoRR, cloned between PstI and HindIII sites, and expressed from pUAB400 in *M. smegmatis*. Lsr2 was amplified by primer pairs FPm2lsr2/RPlsr2 or FPm1lsr2/RPm2lsr2, cloned between PstI and HindIII, or HindIII and ClaI sites of pUAB300, and pUAB400, respectively, and were expressed from the corresponding expression constructs. IdeR was cloned between BamHI/HindIII sites using FPideRstart/RPideRstop primer pair and expressed from pUAB300 (hyg^R^). The co-transformants of *M. smegmatis* were grown on 7H10 plates containing kan and hyg either in the presence or absence of 10 µg/ml Trimethoprim (TRIM) as described earlier (48). In all cases, *phoP*/*phoR* co-expressing constructs, in either combination, was used as a positive control. Oligonucleotide primers and plasmids used to clone and express fusion proteins for M-PFC - based studies are shown in Table S5. The constructs generated in this work were checked by automated DNA sequencing.

### Statistical analysis

Data are presented as arithmetic means of the results obtained from multiple replicate experiments ± standard deviations. Statistical significance was determined by Student’s paired t-test using Microsoft Excel or Graph Pad Prism. Statistical significance was considered at P values of 0.05 (*P≤0.05; **P≤0.01; ***P≤0.001; ****P≤0.0001).

## Acknowledgements

We acknowledge G. Marcela Rodriguez and Issar Smith (PHRI, New Jersey Medical School - UMDNJ) for *phoP*-KO mutant, the complemented mutant strain, and Adrie Steyn (University of Alabama) for pUAB300/pUAB400 plasmids. We thank Punjab Biotechnology Incubator Board for their technical support in ICP-MS studies of our samples, and Christina Shalom Rangarajan for some preliminary results. This study was supported by funding from CSIR-IMTECH intramural grant OLP-0170, CSIR-funded project FBR070308, and SERB-funded project (CRG/2023/00109) to D.S. K.M., H.G., and B.B. were supported by CSIR, and Kajal was supported by DST Inspire fellowship. The funders had no role in study design, data collection and analysis, decision to publish or preparation of the manuscript.

## Author Contributions

K. M., Kajal., B.B., and H. G. designed, performed and analysed the experiments. D.S. conceived and coordinated the study and wrote the paper. All authors reviewed the results and approved the final version of the manuscript.

## Conflict of interest statement

The authors declare that they have no conflict of interests.

## Data availability statement

All the data pertaining to the results reported in this study are either part of the main text or supplemental information files.

